# Pervasive non-CpG methylation at zebrafish mosaic satellite repeats

**DOI:** 10.1101/2020.05.13.093203

**Authors:** Samuel E Ross, Allegra Angeloni, Alex de Mendoza, Ozren Bogdanovic

**Affiliations:** Genomics and Epigenetics Division, Garvan Institute of Medical Research, Sydney, New South Wales, 2010, Australia; St Vincent’s Clinical School, Faculty of Medicine, University of New South Wales, Sydney, New South Wales, 2010, Australia; School of Biological and Chemical Sciences, Queen Mary University of London, London, E1 4NS, United Kingdom; School of Biotechnology and Biomolecular Sciences, University of New South Wales, Sydney, New South Wales, 2052, Australia

**Keywords:** DNA methylation, embryogenesis, zebrafish, repetitive elements

## Abstract

In vertebrates, DNA methylation predominantly occurs at CG dinucleotides even though widespread non-CG methylation (mCH) has been reported in mammalian embryonic and neural cells. Unlike in mammals, where mCH is found enriched at CAC/G trinucleotides and is tissue-restricted, we find that zebrafish embryos as well as adult somatic and germline tissues display robust methylation enrichment at TGCT positions associated with mosaic satellite repeats. These repeats reside in H3K9me3-marked heterochromatin and display mCH reprogramming coincident with zygotic genome activation. Altogether, this work provides insight into a novel form of vertebrate mCH and highlights the substrate diversity of vertebrate DNA methyltransferases.

## Background

Methylation of cytosines within the CG dinucleotide context is the most abundant DNA modification in vertebrate genomes [1]. CG methylation (mCG) is found in all vertebrate cell types and is known to participate in long-term gene silencing processes [2]. Nevertheless, methylation of other cytosine dinucleotides (mCH, H = T,G,A) has also been described. mCH (also referred to as non-CG methylation), is most commonly found in mammalian embryonic stem cells (ESCs) and in the brain, however, mCH has been identified at low levels in almost all human tissues [3–5]. The functions of mCH in gene regulation remain unresolved to date. In mammalian brains, mCH levels are inversely correlated with transcription of the associated gene, whereas this pattern appears to be the opposite in ESCs [4]. mCH within the CA context has also been previously observed at major satellite repeats in mouse ESCs but not in differentiated cells [6]. In mammals, mCH accumulates during nervous system development, and displays considerable remodelling during iPSC reprogramming and direct conversion of fibroblasts to neurons [4,7–9]. Unlike mCG, mCH methylation is not maintained after cell division by DNMT1, and therefore requires constant activity of *de novo* DNMT3 enzymes. mCH deposition is carried out by both DNMT3A and DNMT3B, mostly at CAC or CAG trinucleotides respectively, suggestive of significant sequence specificity during DNMT3 recruitment [10]. Preferential methylation at CAC trinucleotides has also been recently reported as a conserved feature across vertebrates, from lamprey to mammalian brains [11]. However, to date, other forms of mCH have not been identified in vertebrates.

## Results and discussion

To evaluate the presence of non-CG methylation during zebrafish development we reanalyzed whole genome bisulfite sequencing (WGBS) data to obtain genotype-corrected mCH profiles of 80% epiboly (gastrula), 24 <u>h</u>ours <u>p</u>ost <u>f</u>ertilization (hpf, pharyngula), 48hpf (hatching) embryos, and adult brain tissue (bisulfite conversion >99,5%) [12]. All samples showed only a minor elevation of methylation at CA dinucleotides compared to the unmethylated lambda genome spike in control, except for the brain sample which had a moderate 1.5 and 2 fold increase at both CT and CA dinucleotides, respectively (**Fig. 1a**). This is in line with the reported mCA enrichments in the zebrafish forebrain [11]. However, when motif calling was performed on the most highly methylated sites in the CH context, the top enriched sequence was consistently CAT<u>GC</u>TAA, with methylation occurring at the TGCT tetranucleotide (**Fig. 1b and Additional File 1:Figure S1**). Methylation was almost exclusively detected on the strand displaying the 5’-TGCT-’3’ motif (>75%) rather than on its reverse complement (5’-AGCA-3’), suggestive of considerable strand specificity during DNMT targeting as well as lack of symmetry typical of CG methylation (**Additional File 1: Figure S1**). Many of these nucleotides displayed substantial mCH of >10%, particularly at repetitive elements where this motif was found to contain the highest levels of mCH and had a notable increase in methylation at later stages of development and in the brain (**Fig. 1c and Additional File 1:Figure S1**). When the repetitive sites of the genome bearing the CAT<u>GC</u>TAA motif were annotated, we found that more than 65% of these sites are located in MOSAT_DR mosaic satellite repeats (GenBank ID: DP000237.1, **Fig. 1d,e**). Hereafter we refer to this motif as the MOSAT motif. Due to potential mappability issues of repetitive elements, we re-mapped uniquely mapping reads covering MOSAT_DR repeats, tolerating 0 mismatches across the entire read (**Additional File 1:Figure S2 and Additional File 2**). Even under such stringent mapping conditions we observe substantial coverage of MOSAT_DR repeats and confirm the strand bias associated with TGCT methylation (**Additional File 1:Figure S2**). Additionally, to confirm that this form of methylation is not the result of a sequence-specific bias of bisulfite conversion [13], we generated enzymatic methylation sequencing (EM-seq) base-resolution libraries of two biological replicates of 24hpf embryos [14](**Additional File 1:Table S1)**. The top methylated MOSAT motifs identified in either EM-seq or WGBS showed proportionate methylation levels when compared to the orthogonal approach (**Fig. 1f,g and Additional File 1:Figure S3**). We therefore conclude that mCH associated with MOSAT motifs is not due to low bisulfite conversion efficiencies or mappability issues associated with these repetitive regions; MOSAT_DR is an actively CH-methylated satellite repeat in zebrafish.

**Figure 1.**
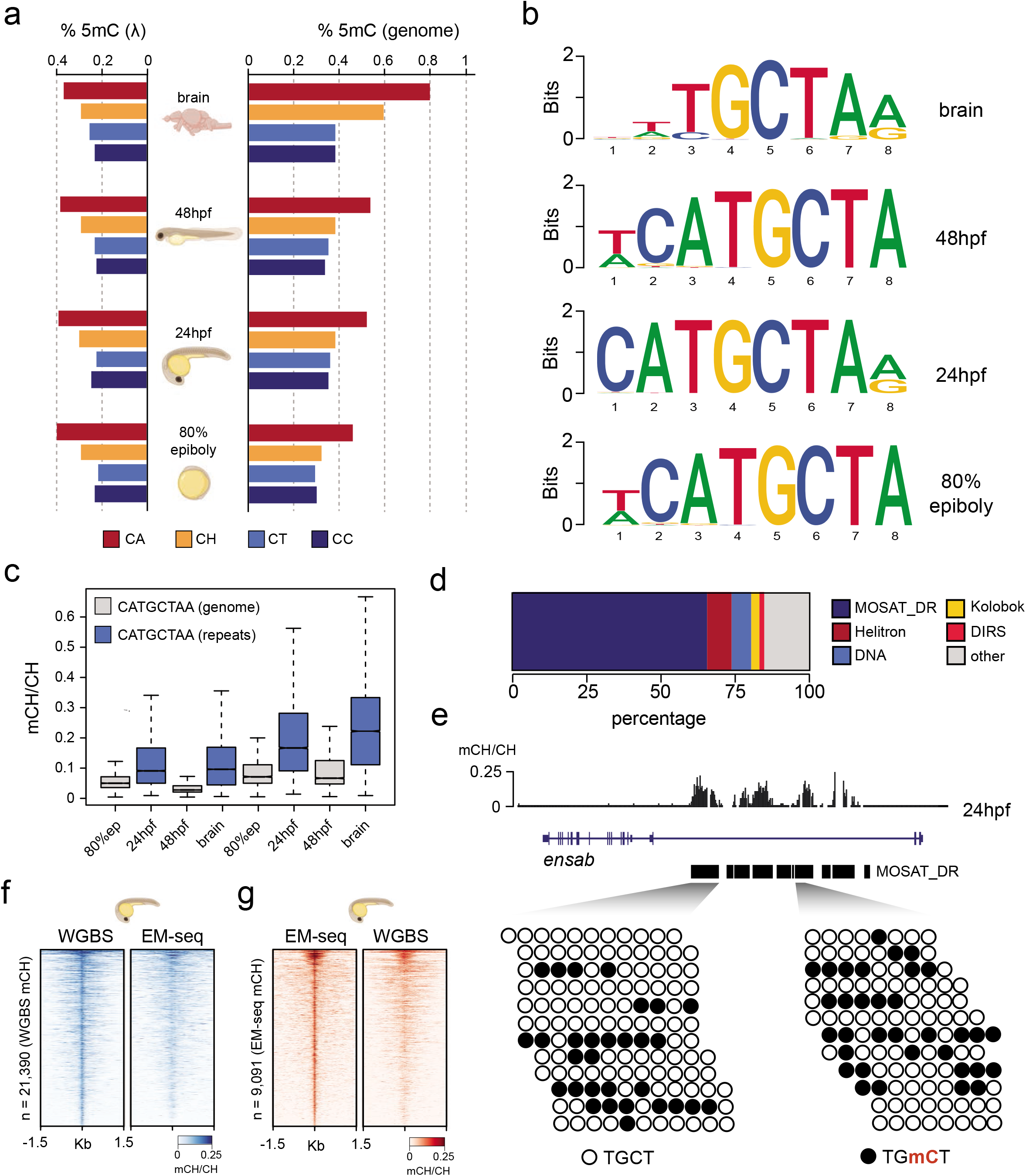
MOSAT motifs are enriched for mCH. **a** Genomic and lambda control dinucleotide %mCH in zebrafish embryos and adult brain. **b** Motif analyses of most methylated CH sites (n=10,000) in zebrafish embryos and adult brain. **c** Boxplots showing average mCH/CH levels of the CATGCTAA motif in repeat-masked and repetitive portions of the genome. **d** Repetitive element annotation of CATGCTAA motifcontaining regions. **e** Per read analysis of MOSAT motif mCH/CH in MOSAT_DR elements. **f** Heatmap of methylated (mCH/CH > 0.1) MOSAT motifs in 24hpf embryos identified by WGBS and compared to EM-seq signal and **(g)** identified by EM-seq and compared to WGBS signal.

MOSAT_DR elements containing the MOSAT motif are found almost exclusively at CG-depleted introns and intergenic regions (**Fig. 2a**). The scarcity of CG dinucleotides in these elements suggests that this type of methylation is unlikely to just be a byproduct of DNMT activity targeting CG sites. In both embryos and adult tissues there is an overall positive correlation between the number of MOSAT motifs and mean mCH levels of the gene observed at all stages, suggesting a major contribution of TGCT methylation to mCH gene body patterning (**Fig. 2b, Additional File 1: Figure S4**). Gene ontology (GO) analysis [15] of MOSAT motif-containing genes demonstrated a significant enrichment for terms associated with neuronal function and in particular synaptic function, in agreement with previous reports on neural mCH enrichment in mammals [4] (**Additional File 1: Figure S4**). Given the observation that long genes can be enriched and sensitive to mCH levels in mice and human brains [16,17], we investigated the length of MOSAT motif-containing genes and found them to be, on average, significantly longer than all genes as well as neural genes (**Additional File 1: Figure S4**). Moreover, MOSAT motif-containing genes do not appear to be constitutively expressed. They exhibit expression in a diverse range of tissues many of which are neural by origin, however, their expression is not exclusively limited to the nervous system [18] (**Additional File 1: Figure S4**).

**Figure 2.**
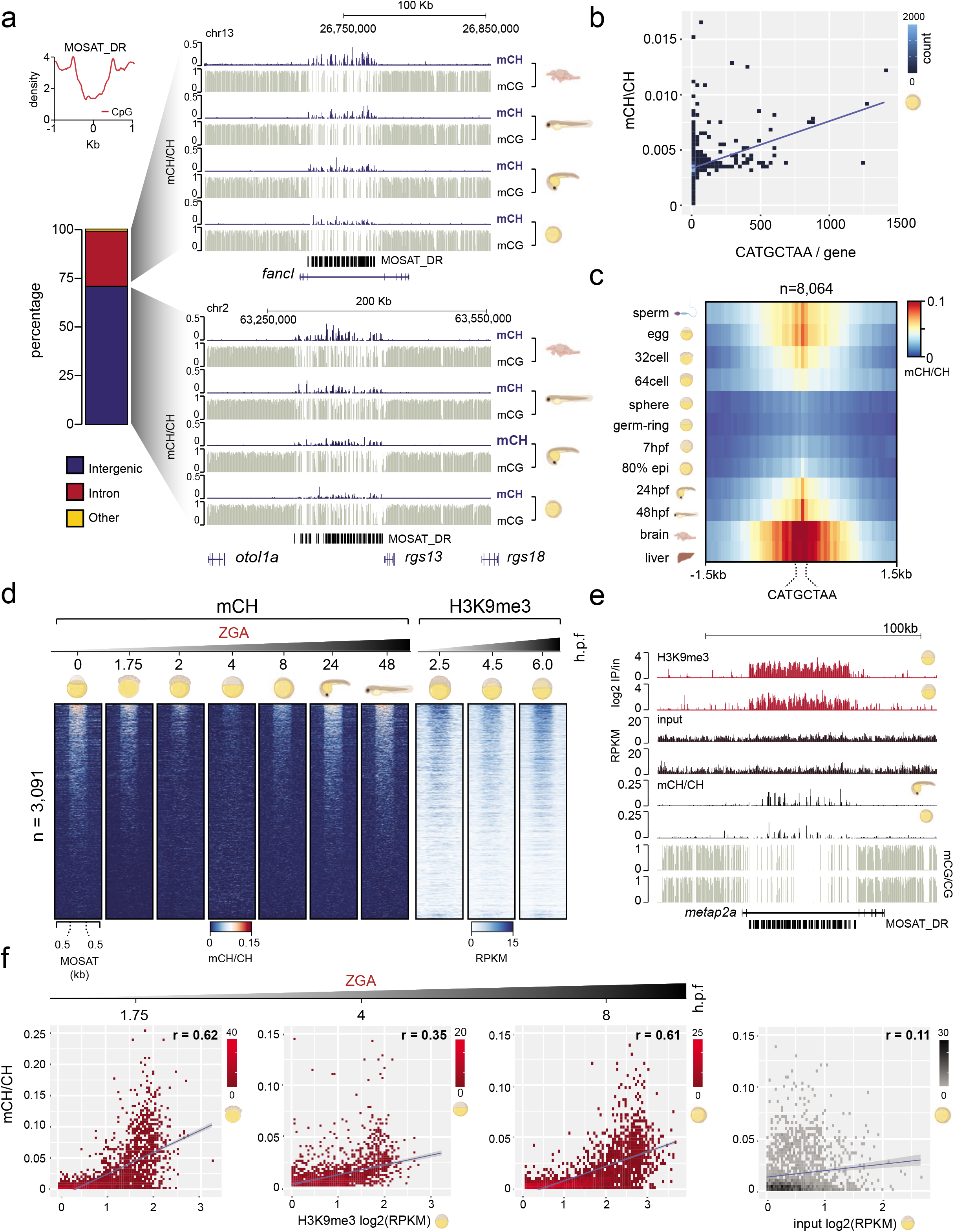
mCH at MOSAT_DR elements is developmentally reprogrammed and associated with constitutive H3K9me3. **a** UCSC browser snapshot of mCH and mCG levels in both intronic and intergenic MOSAT_DR elements in developing zebrafish embryos and adult brain. **B** Scatterplot of average gene methylation (mCH/CH) plotted against the number of MOSAT motifs in gene bodies of 80% epiboly zebrafish embryos. **c** mCH profiles of MOSAT motifs (n=8,064) in commonly covered MOSAT_DR elements during zebrafish development. **d** Heatmap of CH methylation (mCH/CH) and H3K9me3 (RPKM) levels in commonly covered MOSAT_DR elements (n=3,091) during zebrafish development. **e** UCSC browser snapshot of mCG and mCH in 80% epiboly and 24hpf embryos compared to H3K9me3 (log2 ChIP/Input) in shield stage and dome embryos. **f** Scatterplots showing the correlation between mCH/CH at various time points and the level of H3K9me3 (log2RPKM) from shield stage embryos. Left to right: 32cell, sphere, 80% epiboly, 80% epiboly vs input.

Numerous reports have previously described the developmental dynamics of zebrafish CG methylation in somatic and germline tissues [12,19–27]. To further investigate the developmental dynamics of mCH, we reanalyzed base-resolution profiles of liver, sperm, egg, 32 cell stage, 64 cell stage, sphere, germring, and shield embryos [21,24] (**Additional File 1: Table S2**). mCH levels of MOSAT motifs in commonly covered MOSAT_DR elements (8064 sites from 3091 elements) revealed high mCH levels in adult germ cells, cleavage stage embryos and late stage embryos/adult tissues but low levels surrounding the sphere stage, which corresponds to the major wave of zygotic genome activation-ZGA (**Fig. 2 and Additional File 1:Figure S5**). This is in accord with steady state transcript abundance of zebrafish *de novo* DNMT3 enzymes that are low in post-fertilization embryos and higher during later developmental stages, indicating their possible roles in the maintenance of this unique form of methylation [28,29] (**Additional File 1:Figure S5**). The observed temporal mCH dynamics are independent of mCG changes at these regions and are uncoupled from global developmental mCG patterning in zebrafish, which progressively increases from 16 cell stage to match the paternal methylome at the ZGA onset [20,21] (**Additional File 1:Figure S5**). However, the dynamic is similar to what has been observed during mammalian DNA methylome reprogramming, where mCG inherited from male and female germ cells is erased during preimplantation development and re-established during gastrulation in all embryonic layers [30–33]. It is also resemblant of overall mammalian mCH dynamics, which is characterized by gradual dilution of oocyte-inherited mCH during cleavage stages and its gradual re-establishment in the nervous system. [4,31,34].

Finally, we wanted to assess the regulatory environment of mCH-methylated MOSAT motifs. To that end, we analyzed ChIP-seq data of H3K9me3, H3K27me3, H3K36me3, H3K27ac, H3K4me1, and H3K4me3 histone modifications [35–37]. We found a clear enrichment of H3K9me3 at methylated MOSAT_DR elements, whereas no other histone modification displayed any signal at these sites (**Fig.2d-e and Additional File1:Figure S5**). H3K9me3 has been shown to mark the zebrafish genome pre-ZGA and to progressively increase from ZGA onwards [35,38], however, we find that at a sub-population of MOSAT_DR sites, H3K9me3 is stable during development (256 cell to shield) (**Additional File1:Figure S5**). This contrasts the observation in mammals, where H3K9me3 and mCH are generally inversely correlated at CG-poor regions [8,29,39]. To interrogate if H3K9me3 could play a role in recruiting or maintaining mCH at MOSAT repeats, we investigated the correlation between mCH and H3K9me3 through development. We observe a strong positive correlation at the 32 cell stage, 80% epiboly and adult brain (*r* >0.6) where mCH is enriched, and a lower correlation at stages surrounding the ZGA (*r* >0.35), when mCH is being reprogrammed (**Fig. 2f and Additional File 1:Figure S5**). A possible explanation for this correlation could be that mCH is re-established by means of H3K9me3 driven recruitment at these regions. This could be facilitated by DNMT3 enzymes as H3K9me3 can recruit DNMT3 to satellite repeat regions in ESCs [40]. Also, a previous report demonstrated H3K9me3 and DNMT3 cooperation during zebrafish embryogenesis [41]. These findings thus merit further investigation into the possible relationships between heterochromatin and mCH, suggesting that the inverse correlation between mCH and H3K9me3 observed in mammals might not be conserved, or that zebrafish display a lineage-specific innovation.

## Conclusions

Our meta-analysis of mCH in developing zebrafish revealed enrichment of TGCT methylation within mosaic satellite repeats. These motifs were highly methylated in adult germ cells, pre-ZGA embryos, post gastrulation embryos, and adult brain and liver. However, methylation was diminished between the sphere stage and late gastrulation, suggestive of developmental remodeling coincident with ZGA onset thus displaying similarities to mammalian methylome reprogramming. We also found mCH at those sites to be highly correlated with H3K9me3 enrichment during early development in contrast to what has been observed in mammals. In conclusion, we have described a novel type of vertebrate mCH, one with a unique sequence specificity, which undergoes developmental loss and re-establishment at H3K9me3-marked heterochromatin regions.

## Methods

### Zebrafish usage and ethics

Zebrafish work was approved by the Garvan Institute of Medical Research Animal Ethics Committees under AEC approval 17/22. All procedures performed complied with the Australian Code of Practice for Care and Use of Animals for Scientific Purposes. Adult wild type (AB/Tuebingen) *Danio rerio* (zebrafish) were bred in a 1 male:1 female ratio. Embryos were collected 0 hours post-fertilization (hpf) and incubated in 1X E3 medium (0.03% NaCl, 0.005% CaCl2, 0.0013% KCl, 99.9557% H2O, 0.008% H14MgO11S) for 24 hours at 28.5°C.

### Genomic DNA extraction

24hpf embryos were dechorionated using 1mg/mL Pronase (Sigma-Aldrich, St. Louis, MO, USA) diluted in 1X E3 medium, snap-frozen in liquid nitrogen and stored at −80°C prior to DNA extraction. Genomic DNA (gDNA) was extracted from 24hpf embryos using QIAGEN DNeasy Blood & Tissue Kit (QIAGEN) according to manufacturer’s instructions in two biological replicates.

### EM-seq library preparation and sequencing

EM-seq library construction was performed using the NEBNext Enzymatic Methyl-seq Kit (New England BioLabs, Ipswich, MA, USA) according to manufacturer instructions with modifications. 0.02ng of unmethylated lambda phage DNA (Promega, Madison, WI, USA) and 0.0001ng of pUC19 plasmid methylated at 100% of CpG sites (New England BioLabs, Ipswich, MA, USA) were used as spike-in controls to determine the efficiency of APOBEC deamination and TET2 oxidation, respectively. Briefly, 200ng of zebrafish gDNA was sonicated to an average insert size of 300bp. Input DNA concentration was selected according to the optimal input amount as recommended by the manufacturer. Sonicated DNA was end repaired followed by ligation of adapters to DNA overnight using NEXTFLEX Bisulfite-Seq Barcodes (PerkinElmer, Waltham, MA, USA). DNA was treated with TET2 for one hour. Following TET2 oxidation DNA was denatured with 0.1M NaOH then treated with APOBEC for three hours. DNA was PCR amplified (8 cycles). Library concentration was quantified through qPCR using KAPA Library Quantification Kit (Sigma-Aldrich, St. Louis, MO, USA) as per manufacturer’s instructions. 150 pmol of the combined libraries with 15% PhiX spike-in was sequenced on the Illumina HiSeq X platform using 150bp paired-end sequencing (high output mode).

### WGBS and EM-seq data analyses

Bisulfite-converted (WGBS) and APOBEC-converted (EM-seq) [14] sequence reads were trimmed with Trimmomatic (ILLUMINACLIP:TruSeq3-SE.fa:2:30:10 SLIDINGWINDOW:5:20 LEADING:3 TRAILING:3 MINLEN:20) [42], and mapped using WALT (-m 5 -t 20 -N 10000000) [43] onto the bisulfite-converted GRCz11 reference (UCSC) containing λ (WGBS and EM-seq) and pUC19 sequences (EM-seq) added as separate chromosomes. The resulting alignments in BAM format were processed with CGmapTools [43,44] (convert bam2cgmap) to obtain methylation calls. ATCGmap files were parsed to discard the CH positions that showed evidence of a CG position in the reads discordant with the reference genome CH annotation [11]. Genomic data were visualized in UCSC [45] and IGV [46] browsers.

### DNA sequence motif analyses

Genotype-corrected CGmap files were filtered for CH positions covered by at least 10 reads, and sorted by methylation level (mC/C). Top 10,000 positions were then extracted from the reference genome using BEDTools [47] taking the flanking upstream and downstream base pairs (n=5), and preserving the strandedness information. The resulting fasta file was used as input for HOMER “findMotifs.pl” function [48], establishing the search for *de novo* (-S 5) motifs of length 8 (-len 8) with the default scrambled background option. Motifs were visualized using the “ggseqlogo” package in R [49]. The motif matrix (CATGCTAA) was constructed using the seq2profile.pl HOMER function (seq2profile.pl CATGCTAA 0 ets) and the genome-wide motif search was conducted using the scanMotifGenomeWide.pl function (with and without -mask option checked) to uncover CATGCTAA motifs in both repetitive- and non-repetitive DNA.

### mCH level calculation and plotting

DNA methylation (mCH) levels at CATGCTAA motifs were calculated using BEDtools [47] (map function, -o sum) by dividing the sum of reads supporting a methylated CH cytosine with the sum of all reads mapping to that position. mCH levels were plotted using the boxplot function in R (www.r-project.org) (outline = FALSE, notch = TRUE), for positions that had an mCH value > 0. Bedgraphs were generated from the corrected CGmap tools output and converted to bigWig using bedGraphToBigwig script from Kent utils (https://github.com/bowhan/kent/tree/master/src/utils). Heatmaps were generated using deepTools [50] computeMatrix and plotHeatmap functions. For WGBS and EM-seq data comparisons the heatmaps were generated with the following parameters: “computeMatrix reference-point -b 1500 -a 1500 -p 4 -bs 25, --missingDataAsZero” whereas for plotting of mCH levels over MOSAT_DR repeats, we used: “computeMatrix scale-regions -m 650 -b 500 -a 500 -p 4 -bs 25 -p 4” with replacement of NAN values with 0 after the matrix file was generated. For profiles (represented as centered heatmaps) the matrices were generated with computeMatrix (computeMatrix referencepoint --referencePoint center -b 1500 -a 1500 -p 4 -bs 50) and plotProfile functions (--plotType heatmap --yMin 0 --yMax 0.15 --perGroup).

### Assessment of mCH in gene bodies

Zebrafish gene models (ENSEMBL Genes 99, GRCz11) were obtained from www.ensembl.org using the BioMart tool. DNA methylation (mCH) levels in gene bodies were calculated using BEDtools [47] (map function, -o sum) and the number of CATGCTAA motifs in genes was obtained with coverageBed function. Scatterplots of mCH levels and CATGCTAA motif numbers were generated using the geom_bin2d function in ggplot2 ((bins=50) + geom_smooth(method=lm)).

### Repeatmasker track analyses

Repeatmsker track file corresponding to GRCz11 genome reference was downloaded from UCSC. The percentage of repeat subfamilies overlapping CATGCTAA motifs was determined with BEDtools (intersectBed). The genomic annotation of MOSAT_DR motifs was carried out with HOMER [48], annotatePeaks.pl.

### Colocalization of histone modification and mCH

ChIP-seq data sequence reads were trimmed with Trimmomatic (ILLUMINACLIP:TruSeq3-SE.fa:2:30:10 SLIDINGWINDOW:5:20 LEADING:3 TRAILING:3 MINLEN:20) [42], and mapped to GRCz11 genome using bowtie2 default settings [51], allowing multi-mapping reads to align to a single (best) genomic location. The resulting alignments in BAM format were deduplicated using sambamba markdup, default settings [52]. RPKM bigWigs were made using deepTools bamCoverage and reads were centered and extended by 300 base pairs (-e 300 -p 20 --normalizeUsing RPKM --centerReads) [50]. For H3K9me3 data sets, where input data was available, subtraction of input signal was also performed using deepTools bigWigCompare (--operation subtract) before heatmaps were plotted [50]. Heatmaps of histone RPKM levels over MOSAT_DR elements were generated using deepTools computeMatrix (computeMatrix scale-regions -m 650 -b 500 -a 500 -p 4 -bs 25 -p 4), with NAN values replaced with 0 after completion, and were visualized with sorting based on the mCH datasets (plotHeatmap --sortUsingSamples). Data for scatterplots were generated using bedtools map, to determine average mCH levels, and bedtools intersect (-abam), bedtools intersect (-c) and samtools flagstat, to calculate H3K9me3 RPKM [47,53]. Scatterplots were generated using the geom_bin2d function in ggplot2 ((bins=75) + geom_smooth(method=lm) and Pearson’s correlation were determined by the rcorr function in R.

## Author contribution

OB conceived the study. SR, OB, and AdM performed bioinformatic analyses. SR and OB wrote the paper. AA performed EM-seq experiments. All authors contributed to, read, and approved the final manuscript.

## Acknowledgments

We thank Michael Geng for the management and breeding of zebrafish colonies used in EM-seq libraries and Ksenia Skvortsova for input into EM-seq experiments. We thank Ryan Lister for input on mCH data analysis and for critical comments on the manuscript. We thank Sriharsa Pradhan and New England Biolabs for assistance with the EM-seq kit. Images of zebrafish embryos and cell types were created with Biorender.

## Availability of data and materials

Data generated for this submission have been uploaded to the Gene Expression Omnibus (GEO) (https://www.ncbi.nlm.nih.gov/geo) under the record: GSE149416. The following secure token has been created to allow review while this submission remains in private status: cbmlqggctrixzip. WGBS datasets analysed in this study were taken from the following GEO records: %80 epiboly (GSM1662779), 24hpf (GSM1662780), 48hpf (GSM1662781) and adult brain (GSM1662792) (https://www.ncbi.nlm.nih.gov/geo/query/acc.cgi?acc=GSE68087) [12]. Egg (GSM1133392), sperm (GSM1133391), 32cell (GSM1133394), 64cell (GSM1133395) and germring (GSM1133398) (https://www.ncbi.nlm.nih.gov/geo/query/acc.cgi?acc=GSE44075) [21]. 4hpf (GSM3484060, GSM3484068), 7hpf (GSM3484061, GSM3484069) and liver (GSM3505059) (https://www.ncbi.nlm.nih.gov/geo/query/acc.cgi?acc=GSE122722) [24]. ChIP seq datasets analysed in this study were taken from the following GEO records: H3K9me3 at 6hpf (GSM3096185, GSM3096186), 4.5hpf (GSM3096189, GSM3096190) and 2.5hpf (GSM3096193, GSM3096194) (https://www.ncbi.nlm.nih.gov/geo/query/acc.cgi?acc=GSE113086) [35]. H3K27me3 (GSM3165233) and H3K36me3 (GSM3165232) at dome stage (https://www.ncbi.nlm.nih.gov/geo/query/acc.cgi?acc=GSE114954) [37]. H3K27ac (GSM803830), H3K4me1(GSM915193) and H3K4me3 (GSM915190) at dome stage (https://www.ncbi.nlm.nih.gov/geo/query/acc.cgi?acc=GSE32483) [36].

## Supplementary information

**Additional file 1:** Supplementary figures S1–S5 and Supplementary tables S1-S2

**Additional file 2:** MicroBam file for visualisation of reads at four genes covered by MOSAT DR elements.

**Figure S1.**
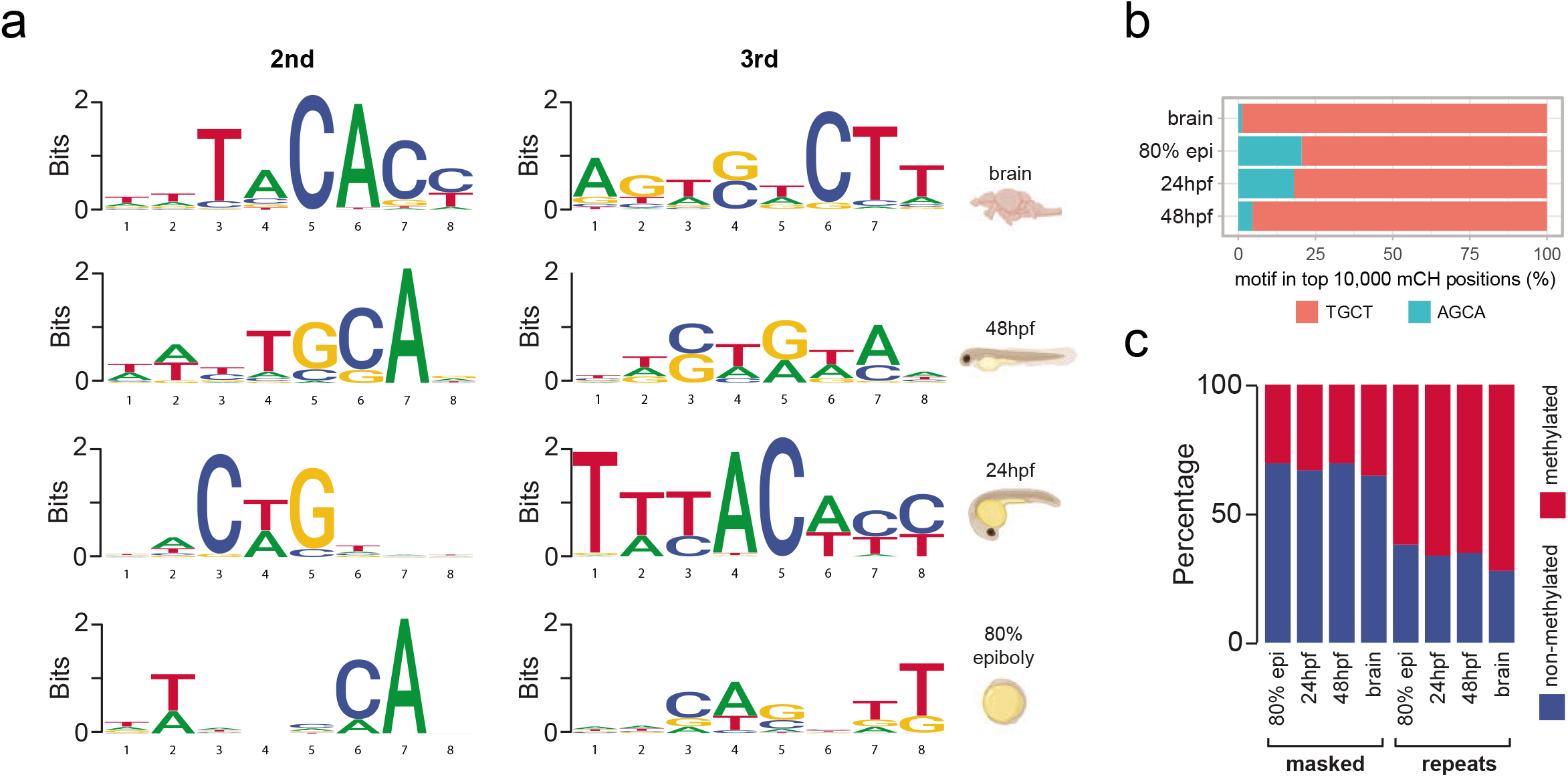
**a** 2nd and 3rd ranked motifs from motif search conducted on 10,000 most highly methyl - ated CH sites in zebrafish embryos and adult brain. **b** Methylation levels of TGCT and AGCA tetranucleotides in 10,000 most highly methylated CH sites in zebrafish embryos and adult brain. **c** Percentage of CATGCTAA motifs exhibiting methylation in zebrafish embryos and adult brains in repetitive and repeat-masked portions of the genome

**Figure S2.**
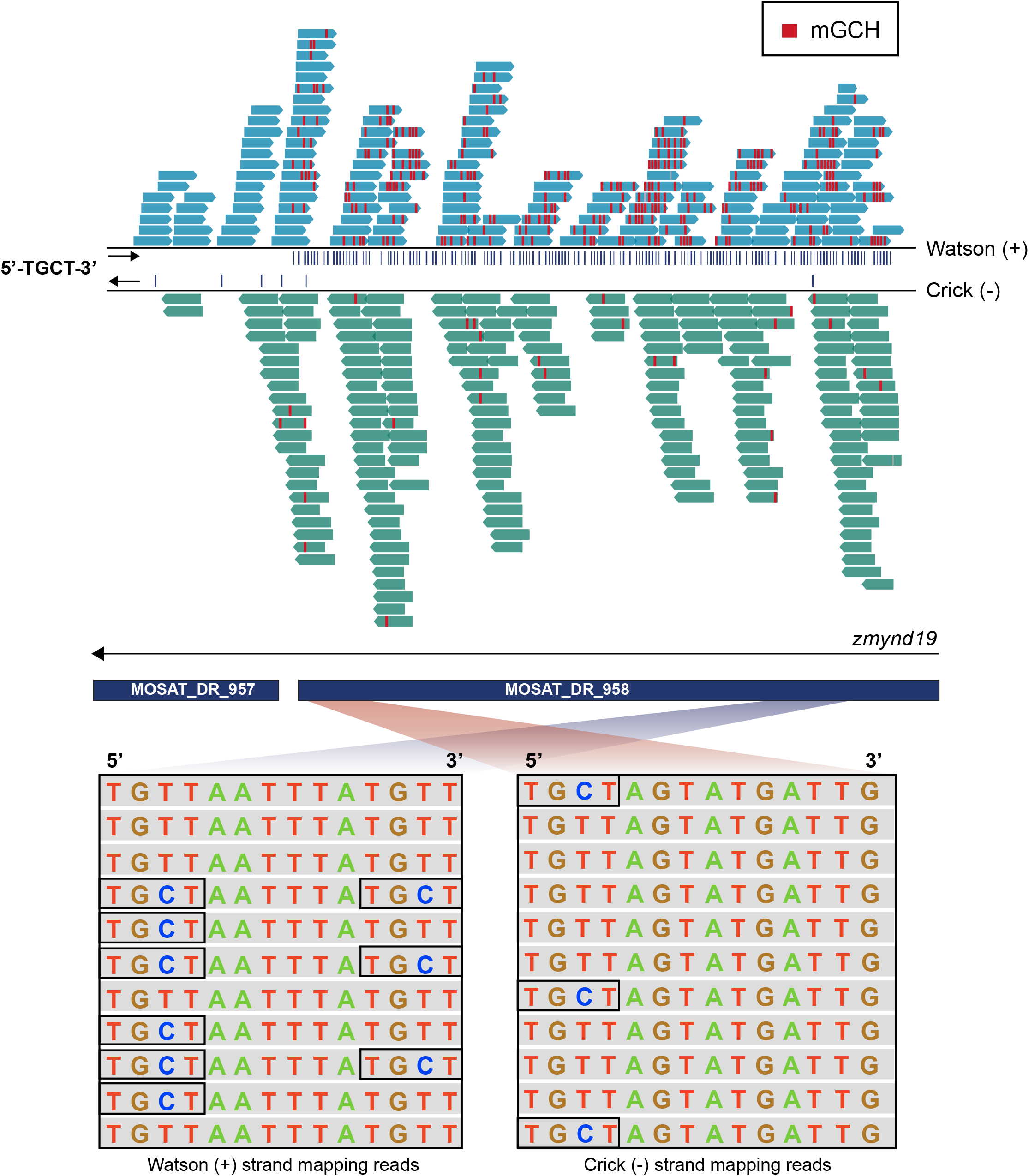
IGV Visualisation of WGBS reads from 24hpf zebrafish embryos mapped with 0 mismatches and aligned to MOSAT_DR elements. Blue reads (forward) map to the Watson(+) strand and contain predominately 5’-TGCT-3’ tetranucleotides (blue arrows) while green reads (reverse) map to the Crick (-) strand. Red lines indicate the positions of methylated GCH nucleotides, which are enriched on the Watson (+) strand and coincide with 5’-TGCT-3’ tetranucleotides. Boxes contain base-resolution visualisation of forward and reverse reads and highlight methylated 5’-TGCT-3’ tetranucleotides within which cytosine did not undergo bisulfite conversion.

**Figure S3.**
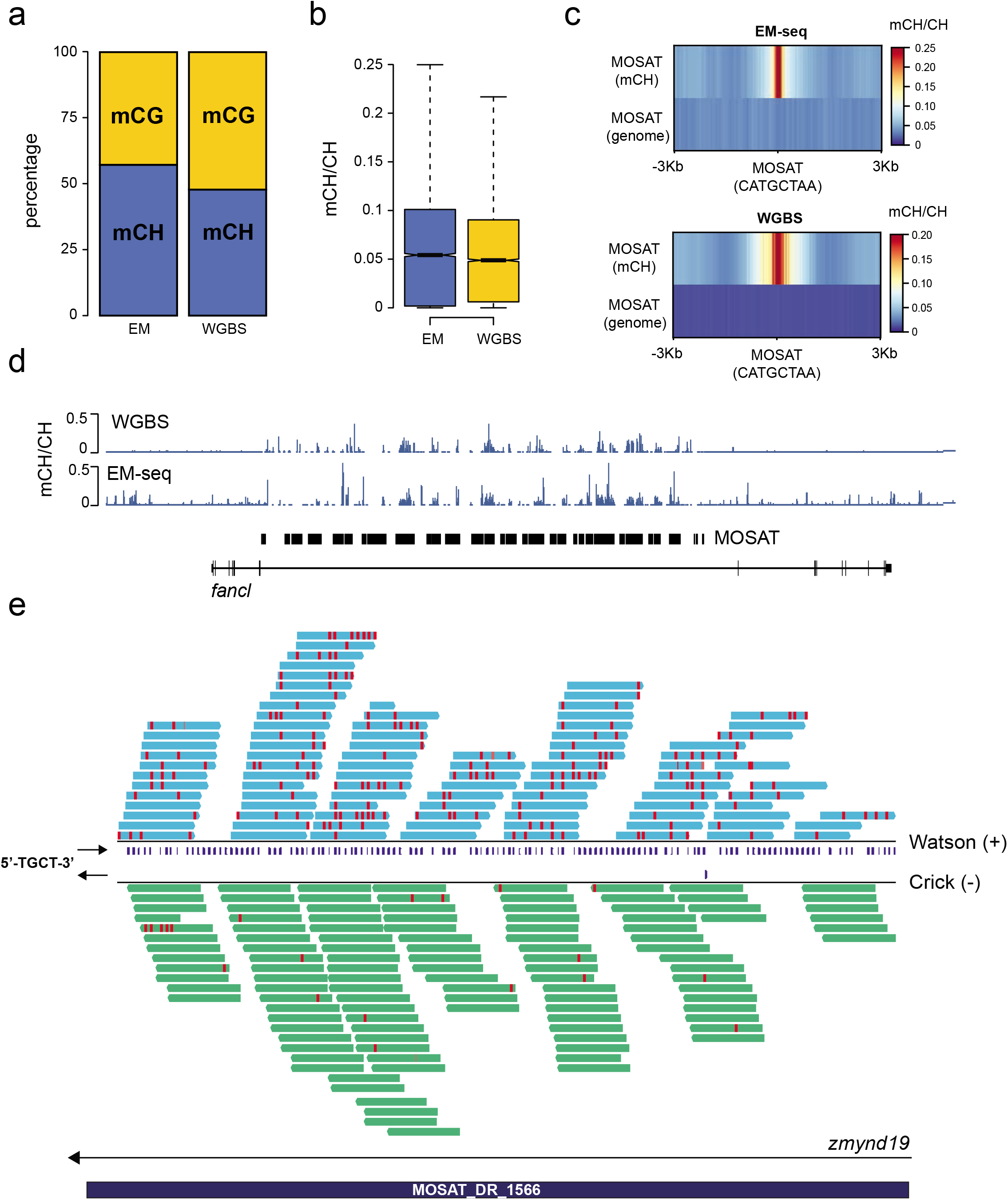
**a** Proportion of methylated sites (≥0.1 mC/C) in MOSAT DR elements in CG and CH contexts determined by EM-seq and WGBS from 24hpf zebrafish embryos. **b** Boxplots of average mCH levels of MOSAT DR elements in 24hpf zebrafish embryos determined by EM-seq and WGBS. **c** Heatmap profiles showing mCH at most highly methylated MOSAT motif sites (≥0.1 mCH/CH) as identified by EM-seq and WGBS in 24hpf zebrafish embryos, and the same number of sites randomly selected from non-repetitive DNA. **d** UCSC browser snaphshot of mCH levels in 24hpf zebrafish embryos at the *fancl* locus, determined by EM-seq and WGBS. **e** IGV Visualisation of EM-seq reads from 24hpf zebrafish embryos aligned to MOSAT_DR elements in the *zmynd19* locus. Blue reads (forward) map to the Watson(+) strand and contain predominately 5’-TGCT-3’ tetranucleotides (blue lines) while green reads (reverse) map to the Crick (-) strand. Red lines indicate methylated GCH positions which are enriched on the Watson (+) strand and 5’-TGCT-3’ tetranucleotides.

**Figure S4.**
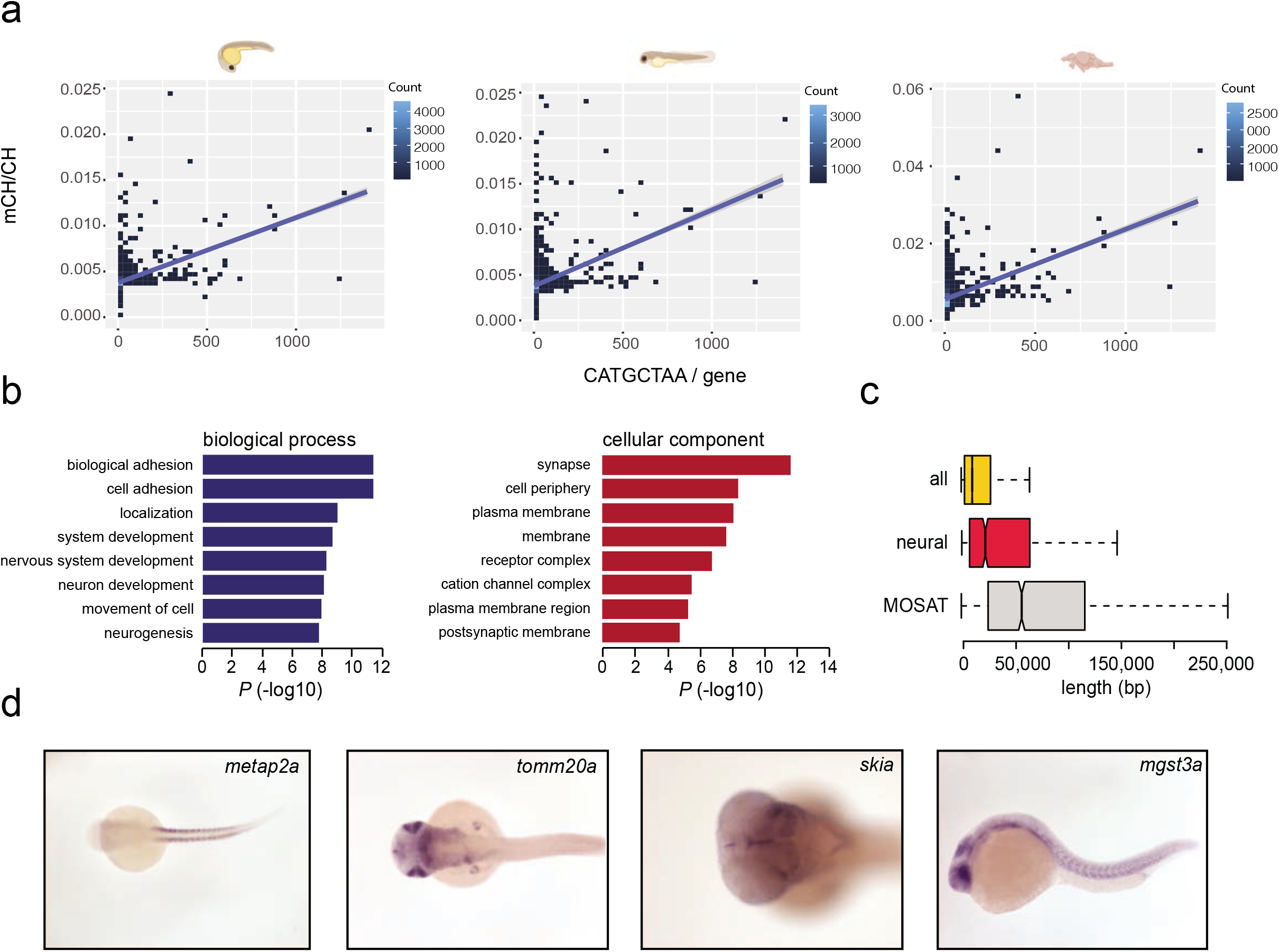
**a** Scatterplots of average gene methylation (mCH/CH) plotted againsy the number of MOSAT motifs in the gene. Left to Right: 24hpf embryo, 48hpf embryo, adult brain. **b** Gene ontology (GO) analysis of MOSAT motif-containing genes. **c** Boxplots of average gene lengths of MOSAT motif-containing genes, neural and all genes**. d** *in situ* hybridisation of MOSAT motif-containing genes (ZFIN database) in zebrafish embryos showing expression of *metap2a* in myotomes (fast muscle fibres), expression of *tomm20a* in retina, tectum, branchial arches and liver, expression of *skia* in the ventral telencephalon, lens, tectum and branchial arches, and expression of *mgst3a* in the eye, tectum and somites.

**Figure S5.**
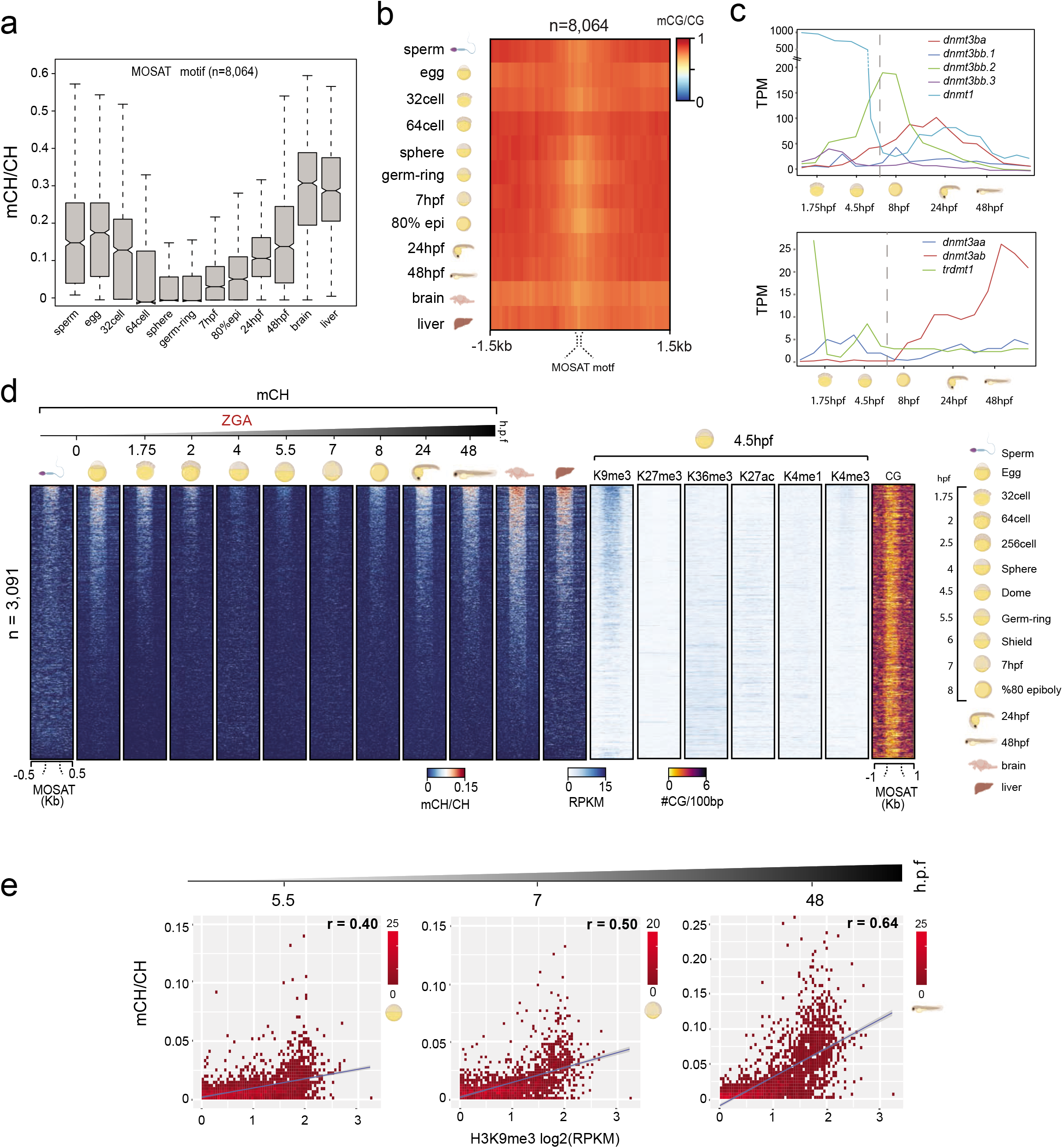
**a** Boxplots of average mCH levels of MOSAT motifs (n=8,066) in developing zebrafish corrected for bisulfte non-conversion rates determined by unmethylated lambda genome spike in controls. **b** mCG profiles of MOSAT motifs (n=8,064) in commonly covered MOSAT_DR elements during zebrafish development. **c** RNA expression analysis of DNMTs during zebrafish development. Dotted lines indicate the estimated time point of mCH re-establishment at MOSAT motifs. **d** Heatmap of mCH levels and histone (RPKM) levels in MOSAT_DR elements (n=3,091) during zebrafish development. **e** Scatterplots showing the correlation between mCH levels at various time points and H3K9me3 at shield stage embryos. Left to right: sphere, 7hpf, 48hpf.

